# Bayesian Analysis of Evolutionary Divergence with Genomic Data Under Diverse Demographic Models

**DOI:** 10.1101/080606

**Authors:** Yujin Chung, Jody Hey

## Abstract

We present a new Bayesian method for estimating demographic and phylogenetic history using population genomic data. Several key innovations are introduced that allow the study of diverse models within an Isolation with Migration framework. For the Markov chain Monte Carlo (MCMC) phase of the analysis, we use a reduced state space, consisting of simple coalescent trees without migration paths, and a simple importance sampling distribution without demography. Migration paths are analytically integrated using a Markov chain as a representation of genealogy. The new method is scalable to a large number of loci with excellent MCMC mixing properties. Once obtained, a single sample of trees is used to calculate the joint posterior density for model parameters under multiple diverse demographic models, without having to repeat MCMC runs. As implemented in the computer program `MIST`, we demonstrate the accuracy, scalability and other advantages of the new method using simulated data and DNA sequences of two common chimpanzee subspecies: *Pan troglodytes troglodytes (P. t.)* and *P. t. verus*.

## Introduction

In the study of diverging populations and species, a common goal is to disentangle the conflicting signals of prolonged genetic drift, which elevates divergence, and gene exchange, which removes it. A widely used conceptual framework for such divergence problems is the Isolation-with-Migration (IM) model, which accounts for genetic drift with parameters for effective population size and splitting time, and for gene exchange with migration rate terms (Fig 1a). IM models have been widely used to study the evolutionary divergence of a very wide range of organisms (Berner *et al.*, 2009; Cong *et al.*, 2015; Geraldes *et al.*, 2008; Hey, 2010a; Moodley *et al.*, 2009; Pinho and Hey, 2010; Won and Hey, 2005).

**FIG. 1.**
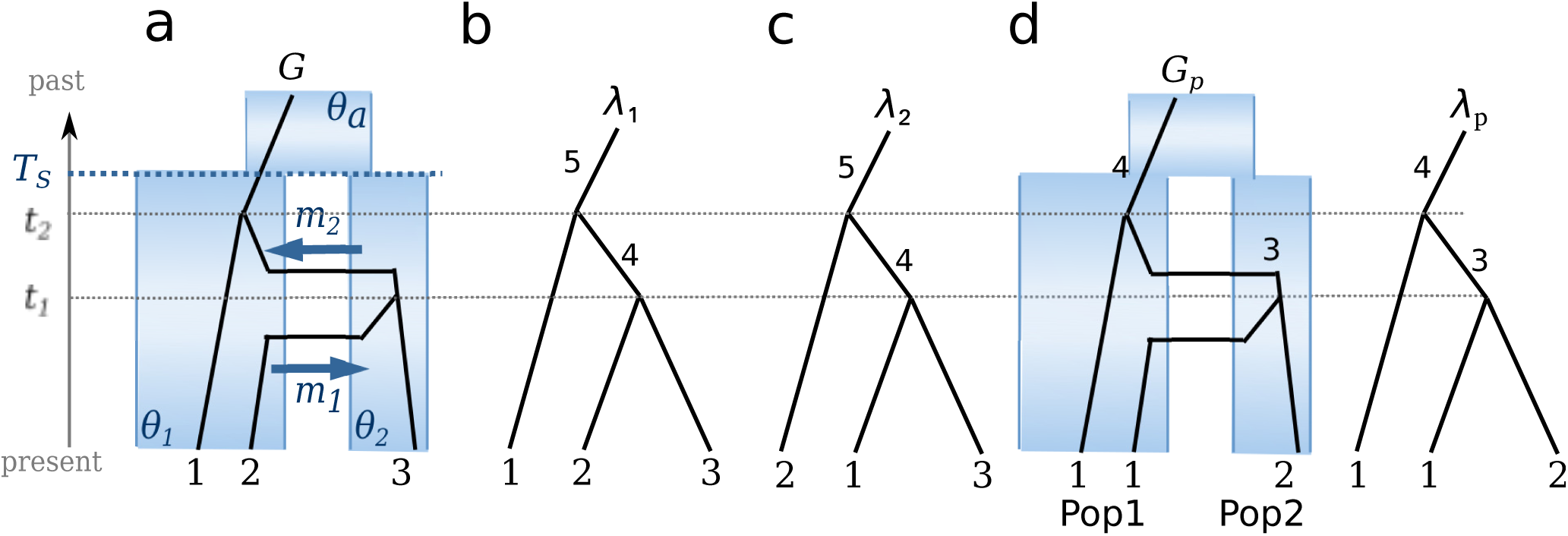
(a) An example of a demographic model with a genealogy. A 2-population isolation with migration (IM) model includes 6 parameters: population sizes, *θ*1,*θ*2 and *θa*, migration rates, *m*_1_ and *m*_2_, and splitting time *T*_*s*_. The graph in black lines depicts a genealogy (*G*) including coalescent events at time *t*_1_ and *t*_2_, respectively, (vertical paths of genes) and migration events (horizontal paths). (b) The coalescent tree (*λ*_1_) of genealogy *G* includes the same coalescent events on *G*. (c) A coalescent tree *λ*_2_ whose probability is same as that of *λ*_1_. (d) Genealogy and coalescent tree with *population labels*.

To connect data, in the form of aligned gene or genome sequences, to the parameters of an IM model, virtually all methods use some form of integration over latent genealogies (Felsenstein, 1988; Griffiths, 1989). A genealogy includes both a coalescent tree, that is an ultrametric binary tree that describes a possible history of common ancestry of a sample of gene copies, and a history of migration events between populations for each of branches in the tree (Beerli and Felsenstein, 1999). Genealogies are not part of the data, nor typically part of the final results, but because we can calculate the probability of aligned sequences given a genealogy (using a mutation model) and because we can calculate the probability of a genealogy given a demographic model (e.g. the parameters for an IM model), likelihood or Bayesian methods for fitting demographic models to aligned DNA sequences all include some kind of machinery for integrating over genealogies (Bahlo and Griffiths, 2000; Griffiths and Tavaré, 1994; Hey and Nielsen, 2007; Kuhner *et al.*, 1995; Lopes *et al.*, 2009; Nielsen, 2000; Nielsen and Wakeley, 2001; Wilson and Balding, 1998).

However inference methods that use IM models frequently face significant computational and statistical challenges. Because of the inclusion of migration events, the space of possible genealogies for a given data set is vastly larger than the space of coalescent trees for the same data. As a practical matter it is difficult to develop a method that adequately samples the space of genealogies, particularly for larger data sets. Likelihood and Bayesian methods for fitting complex demographic models are generally slow and typically cannot be applied to large population genomic data sets (Kuhner, 2008).

Recently, progress has been made on disentangling the migration events from the coalescent tree in the genealogy to allow for calculating the probability distribution of a coalescent tree under an IM model (Andersen *et al.*, 2014; Hobolth *et al.*, 2011; Zhu and Yang, 2012). By representing the history of coalescence and migration using a Markov chain, it becomes possible to integrate over all possible migration histories to calculate the prior probability of a coalescent tree. For example Zhu and Yang (2012) developed a maximum likelihood estimation under an IM model for three DNA sequences using the probability of a coalescent tree.

Here we address several problems associated with genealogy-sampling approaches to demographic inference and present a new Bayesian Markov chain Monte Carlo (MCMC) method for demographic/phylogenetic models including IM models. Our new approach allows for the study of large numbers of loci and can be used for a wide range of demographic models, while allowing for likelihood ratio tests (LRTs) of nested models using a posterior density, proportional to the likelihood, that is joint for all parameters in the model.

To improve the MCMC process and to facilitate the integration over genealogies, we decompose a genealogy *G* into (1) coalescent tree, a simple bifurcating tree, *λ* (Fig 1b), and (2) the remaining information 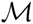, which includes horizontal migration paths of genes between populations. We derive explicit formulas for the probability distribution of a coalescent tree using a Markov chain as a representation of genealogy and matrix exponentiation. To alleviate the computational complexity, we reduce the spaces of genealogies and the state-space of a Markov chain. For efficient MCMC simulations of coalescent trees, we employed importance sampling in which trees are sampled from a tractable probability distribution, rather than from the coalescent probability conditional on the demographic model of interest. Then in the numerical integration over λ, each value of *λ* is weighted by the inverse of its importance sampling distribution. This adjustment accounts for having sampled from the importance sampling distribution and yields an approximation converging to the exact integration over *λ* as more trees are sampled (Robert and Casella, 2013). For the importance sampling distribution, we consider posterior probability distributions in which priors on *λ* are free of the underlying demographic model. Because the coalescent trees do not include migration events and are not constrained by demographic epochs, the MCMC simulation is largely free of mixing difficulties and works well with large numbers of loci.

The computer program, `MIST` (for “model inference from sampled trees”), implements the new method, for multiple processes in parallel. The program `MIST` is freely available at https://github.com/yujin-chung/MIST.git Using simulated DNA sequences, we demonstrate the use of importance sampling distributions and assess the performance of the method in terms of accuracy and computing time. We also demonstrate the application of different models, by using models with and without an unsampled “ghost” population. Moreover, the false positives of LRTs for migration rates are examined based on joint distributions when data show low divergence. Finally, we apply the method to population genomic samples from two subspecies of common chimpanzees (Prado-Martinez *et al.*, 2013), *Pan troglodytes (P. t.) troglodytes* and *P. t. verus*, and compare the results to those from previous other studies.

## New Method

The new method is described for a basic 2-population isolation with migration (IM) model (Fig 1a) with the sizes of two sampling populations and their common ancestoral population (*θ*_1_, *θ*_2_, and *θ_a_*), two migration rates between two sampling populations (*m*_1_ and *m*_2_), and the splitting time of two populations from their common ancestral population (*T*_*S*_). A 2-population IM model with six parameters (Ψ = (*θ*_1_,*θ*_2_,*θ_a_,m*_1_,*m*_2_,*T_S_*)) is easily adapted to variations of this model, such as those shown in Figure 2. Also it should not be difficult to extend the approach to models for data that have been sampled from more than two populations (Hey, 2010b).

Following Felsenstein (1988), the likelihood of Ψ can be obtained by integrating out all possible genealogies in the model:

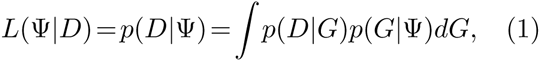

where *p*(*D|G*) is the probability of genetic data *D* given genealogy and *p*(*G*|Ψ) is coalescent probability of genealogy given a demographic model. Considering a Bayesian approach, the posterior distribution of demographic parameters Ψ, given genetic data, is

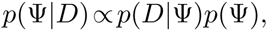

where *p*(Ψ) is a prior distribution under which parameters are independent and uniformly distributed (i.e. *p*(Ψ) is constant over a specified range of values for Ψ).

In considering how to ameliorate the difficulties of working with genealogies it is important to note two things about how a history of migration events impacts the data. First, from a coalescent perspective, the effect of migration events in the true history of a set of genes is to shape the times of common ancestry of those genes. Second, the calculation of the likelihood *p*(*D|G*) depends only on the vertical branch lengths and topology of *G* and not on the migration events in *G*. We can decompose a genealogy into two parts, a coalescent tree *λ* and migration events 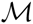, such that 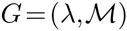 (Fig 1a). Migration will have shaped the coalescent tree, but when the coalescent tree is known, the data are independent of 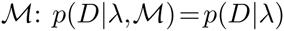. Then using this fact, the integration in Eq (1) separates into two integrations:

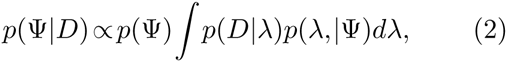

where

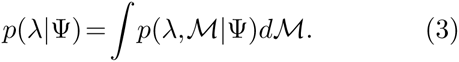

Expression (2) is our target, the posterior density for the model parameters. Below we provide a formula for computing analytically the integration over 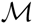, in Eq (3) and we develop a new MCMC approach using importance sampling to approximate the integration over *λ* in Eq (2).

### Exact integration over migrations: Markov chain representation

The exact integration over migration paths in Eq (3) is done by computing transition probabilities in a Markov chain representation of a genealogy *G*. To reduce the state space of the Markov chain representation of a genealogy *G*, we introduce a simplified genealogy in which sampled gene copies are labeled only by the *label* of the population they were sampled from. We define a function *ϱ* that replaces the tip labels on *G* or *λ* by the label of their respective sampled population. The genealogy or coalescent tree with *population labels* is denoted as *ϱ*(*G*) = *G*_*p*_ or *ϱ*(*λ*) = *λ*_*p*_, respectively. For example, *λ*_1_ and *λ*_2_ in Figure 1 have different individual tip labels, but are converted to the same coalescent tree with *population labels*: *ϱ*(*λ*_1_) = *ϱ*(*λ*_2_) = *λ*_*p*_. Moreover, the probabilities of *λ*_1_ and *λ*_2_ are the same: *p*(*λ*_1_|Ψ) = *p*(*λ*_2_|Ψ). In general, any set of trees that can be converted into the same coalescent tree using population labels, will share the same probabilities (see Lemma in the Supplementary Information). Using this property and the following Theorem 1, we compute *p*(*λ*|Ψ) from *p*(*λ_p_*|Ψ).

#### Theorem 1.

*Consider a m-population IM model with parameters* Ψ. *Let* Λ_*p*_ = {*λ|ϱ*(*λ*) = *λ_p_} be the collection of coalescent trees that are converted into the same coalescent tree with population labels λ_p_. Then its size is*

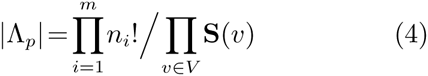

*where V is the set of vertices of λ_p_ that has two tips as descendants (so called “cherry”), n_i_ is the number of samples from population i (i* = 1,*…,m) and* **S**(*v*) *is 2 if the two descendants (tips) of v have the same labels; 1 otherwise. The probability density of λ ∈* Λ_*p*_ is

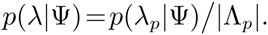

The proof of Theorem 1 is provided in the Supplementary Information. Depending on the sample configuration this can considerably reduce the total number of trees used in the computations.

To compute *p*(*λ_p_*|Ψ) we used a Markov chain representation of *G*_*p*_ in which time is separated into multiple epochs bounded by coalescent times and population splitting times. For example, in Figure 1d, two epochs, (0,*t*_1_] and (*t*_1_,*t*_2_], are defined by two coalescent events at time *t*_1_ and *t*_2_, respectively. The genealogy in each time epoch can be expressed as a sequence of transitions (Asmussen, 2003) among transient states (migration events) and into absorbing states (coalescent events). A state *s* of a Markov chain {*X*(*t*)} is a subset of {(*l,q*): *a|a ≥* 0,*l* = 1,*…,k*;*q* = 1,*…,p*}, where *a* in (*l,q*): *a* denotes the number of lineages with label *l* in population *q* and *k* is the total number of kinds of lineages’ labels. Note that tips on *G*_*p*_ or *λ*_*p*_ may have the same labels, but ancestral lineages have distinct labels. In Figure 1, the lineages on genealogy *G*_*p*_ have labels 1 to 4 and all transient states in each epoch are in Table 1. The initial state of *G*_*p*_ at time 0 is *s*_2_, the state right before the first coalescent event at time *t*_1_ is *s*_4_, and the state right after the event is 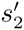. If a state has an element (*l,q*): 0 of zero number of lineages for some *l* and *q*, then, for an efficient expression, we consider the states with and without the element with no lineage are identical. For example, *s*_1_ = {(1,1): 2,(2,1): 1} = {(1,1): 2,(2,1): 1,(1,2): 0}.

**Table 1.** The possible transient states of Markov chains as representatives of *G*_*p*_ in epoch (0,*t*1] and (*t*1,*t*2], respectively.

| | Epoch 1 $(0, t_1]$ | | Epoch 2 $(t_1, t_2]$ |
| --- | --- | --- | --- |
| $s_1$ | $\{(1,1):2,(2,1):1\}$ | $s'_1$ | $\{(1,1):1,(3,1):1\}$ |
| $s_2$ | $\{(1,1):2,(2,2):1\}$ | $s'_2$ | $\{(1,1):1,(3,2):1\}$ |
| $s_3$ | $\{(1,1):1,(2,1):1,(1,2):1\}$ | $s'_3$ | $\{(1,2):1,(3,1):1\}$ |
| $s_4$ | $\{(1,1):1,(1,2):1,(2,2):1\}$ | $s'_4$ | $\{(1,2):1,(3,2):1\}$ |
| $s_5$ | $\{(1,2):2,(2,1):1\}$ | | |
| $s_6$ | $\{(1,2):2,(2,2):1\}$ | | |

In general, the transition rate *q*_*i,j*_ from state *s*_*i*_ to state *s*_*j*_ is as follows:

1. if *s*_*i*_\*s*_*j*_ = {(*l,p*): *a,*(*l,q*): *b*}, *s*_*j*_\*s*_*i*_ = {(*l,p*): (*a−*1),(*l,q*): (*b*+1)} (i.e., a lineage with label *l* moves from population *p* to *q*), then *q*_*i,j*_ = *am*_*p,q*_, where *m*_*p,q*_ is the migration rate from population *p* to population *q* backward in time, and the set difference *v \w* is defined by *v \w* = {*x ∈ v | x* ∉ *w*}.
2. if *s*_*j*_ = *A* (the absorbing state), *X*_1_ = {(*a,p,l*)|(*l,p*): *a ∈ s_i_,a ≥* 2}, *X*_2_ = {(*a,b,p,l,l′*)|(*l,p*): *a∈s_i_,*(*l′,p*): *b∈s_i_,a ≥* 1,*b ≥* 1,*l > l′*}, and either *X*_1_ or *X*_2_ is not an empty set (i.e., two lineages with the same label *l* or different labels *l* and *l′* coalesce),

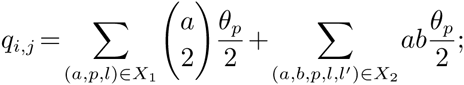
3. if *i* = *j*, then *q*_*i,j*_ = *−* ∑_*j≠i*_ *q*_*i,j*_;
4. otherwise, *q*_*i,j*_ = 0.

For example, the state change from *s*_1_ = {(1,1): 2,(2,1): 1} = {(1,1): 2,(2,1): 1,(1,2): 0} to *s*_3_ in Table 1 means that a lineage with label 1 migrates from population 1 to 2. The transition rate for the event is *q*_1,3_ = 2*m*_1_, since *s*_1_ *\s*_3_ = {(1,1): 2,(1,2): 0} and *s*_3_ *\s*_1_ = {(1,1): 1,(1,2): 1}. Similarly, the formulas are applied for every transition event. The transition rate matrices *Q*_1_ and *Q*_2_ for *G*_*p*_ in epoch (0,*t*_1_] and (*t*_1_,*t*_2_], respectively, are below:

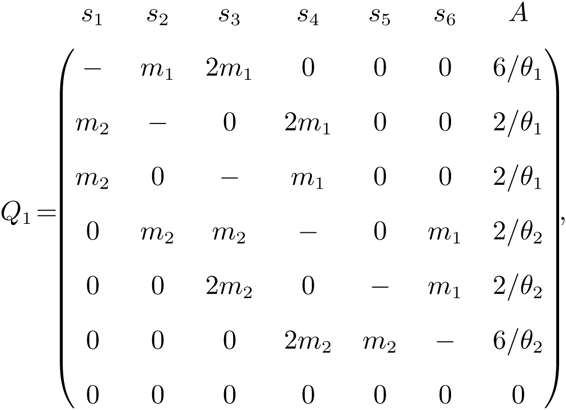

and

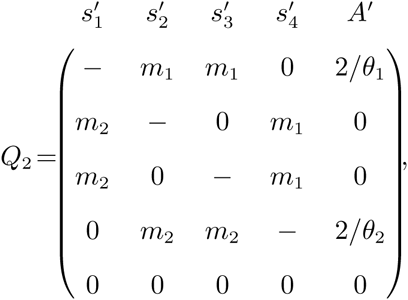

where diagonal elements are set to the negative sum of the corresponding row. Then the probability of state change from *s*_*i*_ to *s*_*j*_ during time *t*_1_ is *P r*(*X*(*t*_1_) = *s_j_|X*(0) = *s*_*i*_) = (*e*^*t*_1_*Q*_1_^)_*i,j*_, where *e*^*Q*^ is a matrix exponential and (*Q*)_*i,j*_ is (*i,j*) entry of matrix *Q*.

Because the coalescent tree *λ*_*p*_ does not include the information in which populations the coalescent events occurred, computing *p*(*λ*_*p*_) requires that we consider all the possibilities. The possible states right before the first coalescent event are *s*_1_ (all lineages in Population 1), *s*_3_ (the coalescing lineages only in Population 1), *s*_4_ (the coalescing lineages only in Population 2) and *s*_6_ (all lineages in Population 2). The corresponding states right after the event are 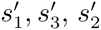, and 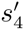, respectively. The probability of the case that *s*_4_ is the state right before the first event is 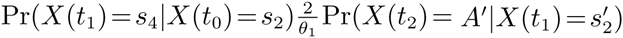. In a similar way, we compute the probability of each possible way and the probability of *λ*_*p*_ takes account of all possible cases:

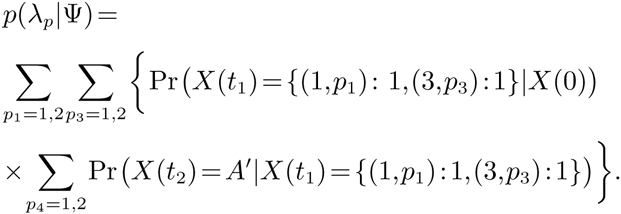

Using this approach we can compute the probability of any coalescent tree under an IM model.

### IM model estimation based on importance sampling of trees

Importance sampling is a widely used approach for working with a distribution of interest, but that for some reason may present difficulties, by using another distribution that is more tractable (Robert and Casella, 2013). In our case we desire coalescent trees sampled from Eq (3) using MCMC, but using an importance sampling distribution to simplify the process. If *q* is a probability density from which we can generate trees easily (i.e. our *importance sampling distribution*) then we can write

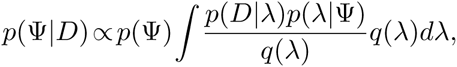

where *q*(*λ*) > 0 if *p*(*D|λ*)*p*(*λ*|Ψ) *>* 0. The above integration can be estimated using *n* draws *λ*_1_,…,*λn* from *q*(*λ*) by the expression,

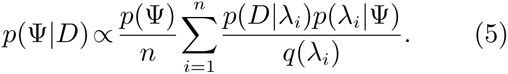

**FIG. 2.**
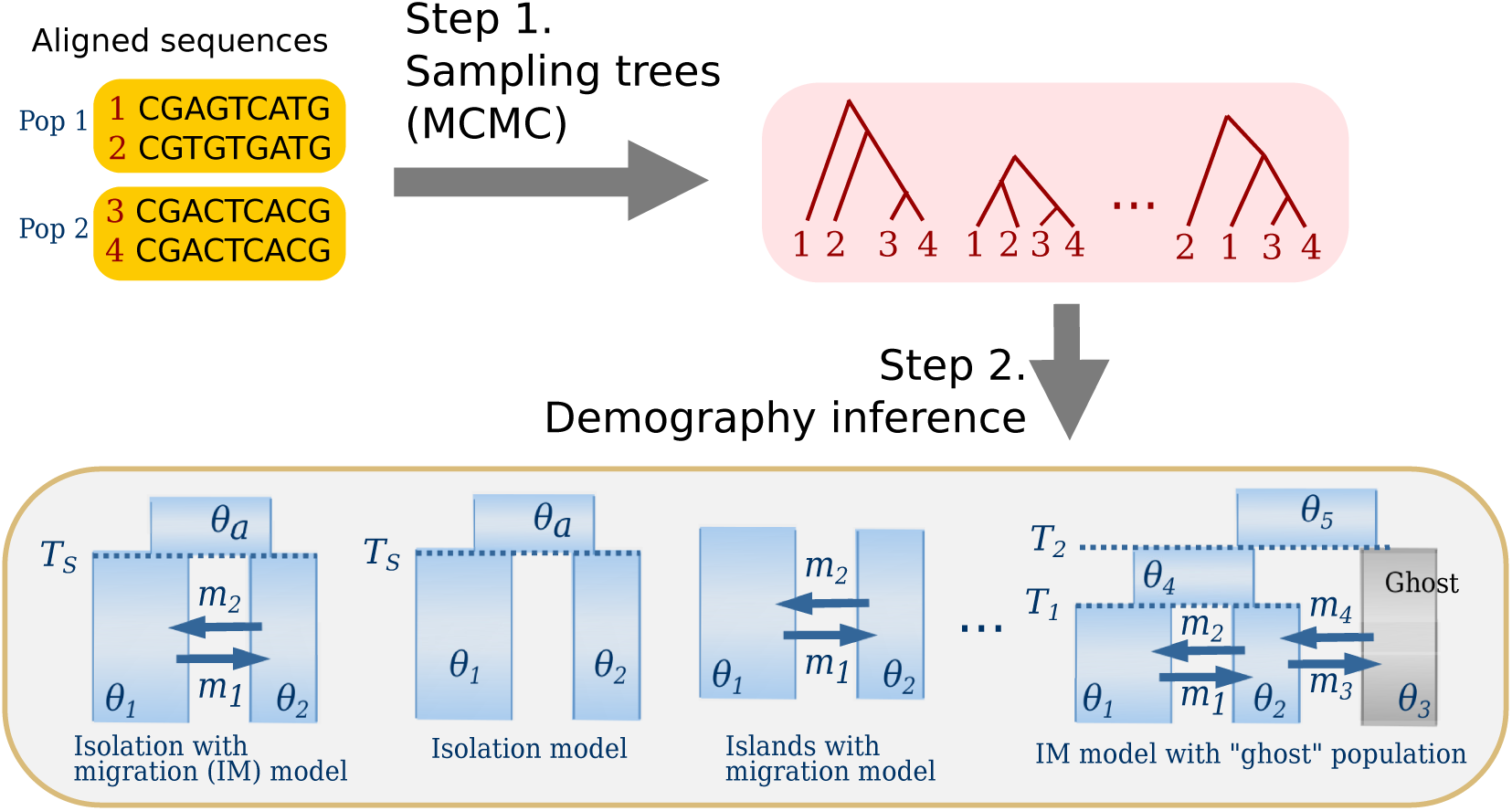
The new method schematic. In step 1 coalescent trees for the aligned DNA sequences are sampled from an MCMC simulation using an importance sampling prior distribution of trees that is free of a demographic model. In step 2, the set of sampled coalescent trees is used for the approximation of the joint posterior density under a demographic model of interest. Optimization of the joint posterior density provides an estimate of model parameters. The same set of trees from step 1 can be used repeatedly to study different demographic models.

We consider two distinct posterior distributions as importance functions, neither of which depends on an underlying demographic model. The first assumes a uniform improper prior, *q*_1_(*λ*) *∝* 1, on the space of coalescent trees, which consists of a finite set of tree topologies and an infinite set of each of branch lengths. This prior is non-informative and does not assume any demographic model. It follows that the sampled coalescent trees are drawn from a posterior distribution that is strictly proportional to the likelihood of the DNA sequences: *q*_1_(*λ|D*) *∝ p*(*D|λ*)*q*_1_(*λ*) *∝ P* (*D|λ*). With the infinite-site model to calculate the likelihood, the posterior density is a proper probability distribution and explicitly a mixture of the product of gamma distributions (see the Supplementary Information). When this importance function *q*_1_(*λ|D*) is applied, the ratio *p*(*D|λ*)*p*(*λ*|Ψ)/*q*__1__(*λ|D*) is proportional to *p*(λ|Ψ). Therefore, with a sample of *λ*_1_,*…,λ_n_ ∼ q*_1_(*λ*|Ψ), the approximation (5) is simplified as

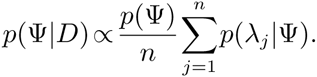

By using this improper prior, we sample coalescent trees mostly where the likelihood is large and hence we expect this importance sampler to be efficient. The second importance function we consider assumes a simple single population model for which the single population size parameter, *θ*, is integrated out analytically. The explicit form of the prior *q*_2_(*λ*) is

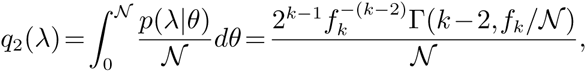

where 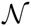 is a constant, *k* is the number of tips on *λ*, 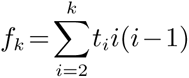 for *t*__2__,*…,t__k__*, coalescent times on *λ* and 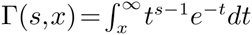, the upper incomplete gamma function. With this importance function, *q*_2_(*λ|D*) *∝ p*(*D|λ*)*q*_2_(*λ*), the posterior density in (5) can be approximated as follows:

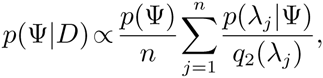

where *λ*_1_,*…,λ_n_ ∼ q*_2_(*λ|D*).

In overview our new approach can be envisioned as a 2-step procedure (Figure 2). In step 1, coalescent trees are sampled from an MCMC simulation under an importance sampling distribution, independent of a demographic model of interest, while in step 2 the joint posterior probability of the demographic model of interest is calculated. Once the sample of coalescent trees has been obtained, they are used to build a function for the joint posterior density of the demographic model of interest, which in turn is used to find the maximum *a posteriori* (MAP) estimate of the model parameters. We used a differential evolution algorithm (Price *et al.*, 2005) to maximize the joint posterior density, but other methods can be used. By using a demography-free importance sampling distribution in step 1, it is possible to study diverse demographic models without having to repeat step 1. For example, with data from two populations, the same coalescent trees sampled in step 1 can be used to examine the data under an IM model, an isolation model, an islands with migration model and an IM model with an unsampled “ghost” population (Fig 2). Another benefit is that by analytically integrating over all possible migration paths in step 2, sampling variance of migration paths is not a source of variance in parameter estimation or model choice as it is in methods that sample migration paths from a MCMC simulation.

### Multiple loci and parameterization of mutation rates

We consider two parameterizations of the mutation process, one in which all loci experience the same mutation rate per site, and a second model in which each locus has its own mutation rate. The constant mutation rate model, in which the mutation rate experience by a locus is proportional to its length, is quite straightforward to implement in the MCMC sampling, even for very large numbers of loci. Under this constant mutation rate model, demographic parameters and coalescent times are scaled by the mean of per-site mutation rate. Therefore, the mutation rate is not estimated through an MCMC simulation and the approximated posterior density is as follows:

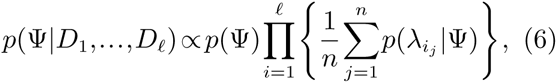

where *λ*_*i,j*_ is the *j*th sampled tree for locus *i* from importance sampling function *q*. Although *n* coalescent trees are given for each locus, the approximation is computed from *n*^ℓ^ joint samples of coalescent trees for *ℓ* loci.

Under the locus-specific mutation rate model, each locus has a mutation rate scalar and the product of all mutation scalars is constrained to be 1, with demographic parameters scaled by the geometric mean of the mutation rates across loci (Hey and Nielsen, 2004). Under this approach the posterior density is approximated as

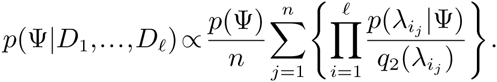

Compared to the constant mutation rate model, the locus-specific mutation rate model includes mutation scalars in the Markov chain state space and requires more iterations in MCMC simulation. The posterior surface to explore in MCMC simulation is the joint probability of mutation scalars and coalescent trees. With many loci, the joint surface would be more difficult to explore, whereas in the constant mutation rate model the number of loci does not affect the number of MCMC iterations.

## Results

We evaluated the performance of the new method using computer simulations. We used *ms* (Hudson, 2002) to simulate 2 gene copies from each of 2- populations IM model with *θ*_1_ = 5, *θ*_2_ = 1, *θ*_3_ = 3, *m*_1_ = 0.02, *m*_2_ = 0.1 and *T*_*S*_ = 2 (Figure 1a), and varied the number of loci, including 10, 100, 1,000 and 10,000. For each case, 20 replicates were generated. We assumed an infinite sites mutation model (Kimura, 1969) and no recombination within loci, but free recombination between loci. We also assumed that all loci have the same mutation rate. For each analysis we sampled 1,000 coalescent trees per locus after 100,000 burn-in iterations and 100 thinning iterations from the MCMC simulation. Convergence diagnostics were monitored (Supplementary Notes and Supplementary Fig 1-2). In step 2, the upper bounds of population sizes, migration rates and splitting time were 20, 10 and 10, respectively. The optimization of the joint posterior density yielded joint MAP estimates for all 6 model parameters.

As shown in Figure 3, the method provides consistent and asymptotically unbiased estimations. The mean of each parameter estimate became closer to the true value (the absolute bias ranges between 0.009 and 0.016 on 10,000-locus data) and the standard errors (range: 0.0028-0.0699 on 10,000-locus data) were also substantially reduced as the number of loci increases. The mean squared errors (MSEs) consisting of bias and variance of estimators were strictly decreasing with the number of loci (Supplementary Table 1). The overall accuracy of all parameter estimations was quite high with just 100 loci, and estimates were very close to the true values with 1,000 or more loci.

We also assessed the performance of the new method in terms of computing time. At each iteration of MCMC simulation in step 1, one coalescent tree of 4 gene copies from each locus is simulated. Thus, the CPU time in serial computing of each iteration is proportional to the number of loci (Figure 4a). In step 2, when the posterior probability of a given demographic model is approximated from a set of coalescent trees, a matrix decomposition is required to compute the probability of a coalescent tree. To avoid repeated computation, matrix decomposition is done for each ranked tree topology with *population labels*. In this simulation study, there are 7 possible ranked tree topology with *population labels* when 2 gene copies were sampled from each population. Then we need to do matrix decomposition for these trees no matter how many loci are analyzed and how many trees are sampled from an MCMC simulation. In our analyses, the computing time of matrix decomposition was constant as 0.02 seconds for the case of 4 sequences. Given the result of matrix decomposition, the CPU time of computing the posterior probability is proportional to the number of loci in a serial computation (Figure 4b). In parallel computing, the computing time of both steps is substantially reduced (Supplementary Table 2) and this method is appropriate to analyze many loci.

**FIG. 3.**
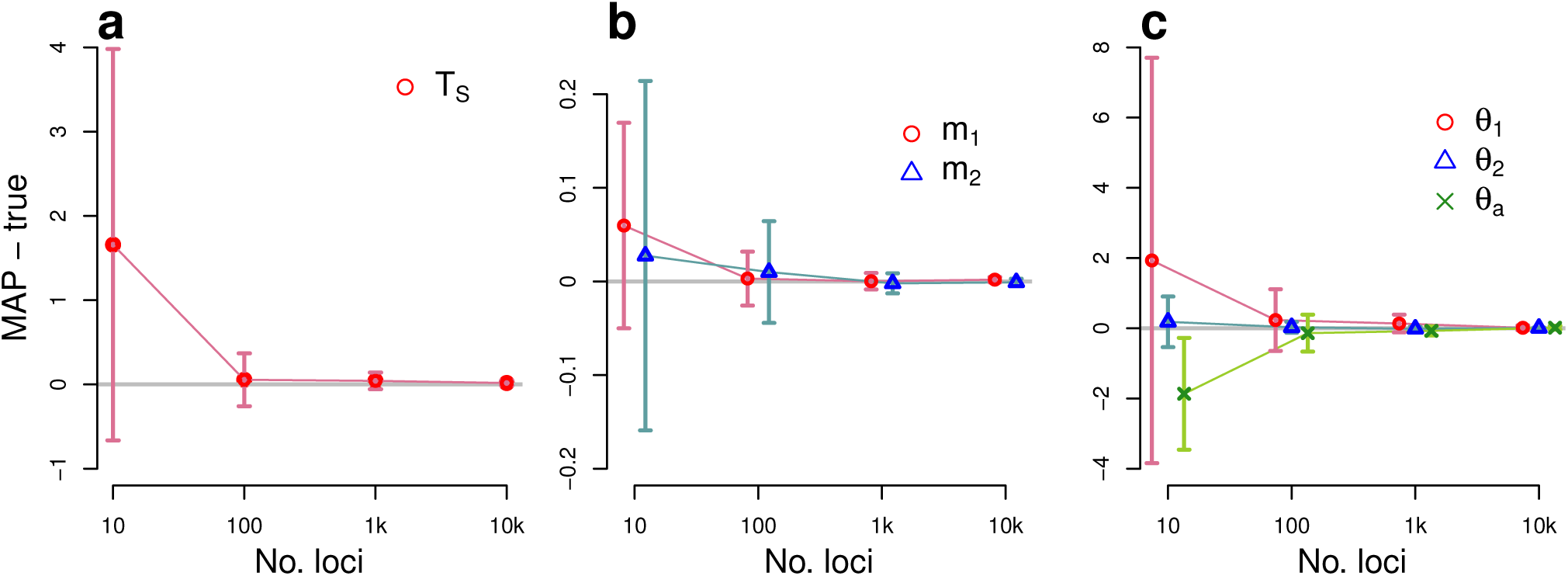
Simulation results illustrating the performance of the new method for a 2-population IM model. The true IM model has parameters *T*_*S*_ = 2, *m*_1_ = 0.02, *m*_2_ = 0.1, *θ*_1_ = 5, *θ*_2_ = 1 and *θ*_*a*_ = 3. DNA sequences were simulated over a range of loci numbers. For each plot, the *x* axis for numbers of loci is on a log scale. The difference between the true value and the mean of the estimated values are plotted (gray horizontal line at 0), and vertical dashed lines indicate standard errors. The average of MAP estimations from 20 replicates, each with 1,000 coalescent trees per locus sampled in step 1, are compared with the true parameters. (a) The average difference between MAP estimates and the true splitting time (b) The average differences for migration parameters. (c) The average differences for the population size parameters for sampled and ancestral populations.

**FIG. 4.**
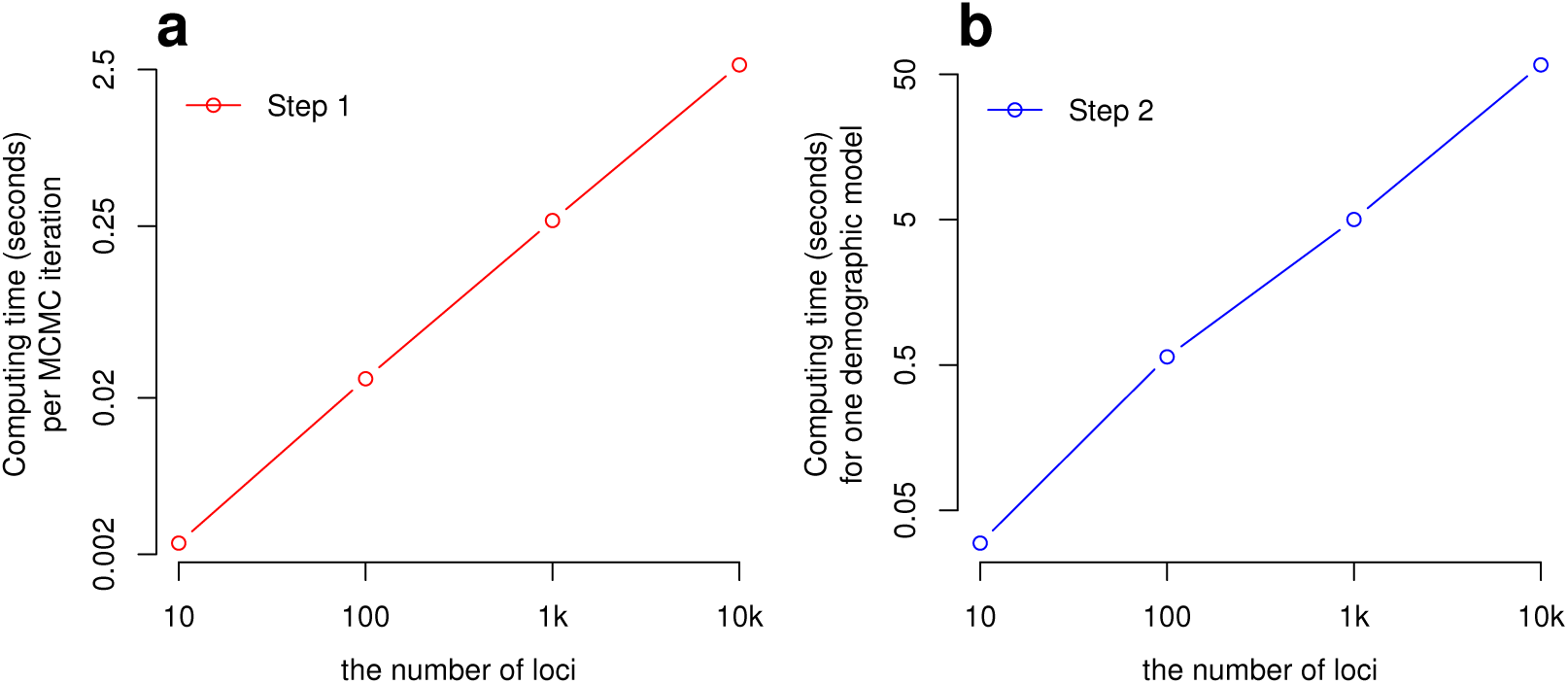
The average CPU time for a single CPU (serial computation) in Step 1 and Step 2 of the new method as a function of the number of loci with 4 gene copies. (a) The mean CPU time for one iteration of MCMC simulation (Step 1), including proposal and evaluation of an update for each locus. Both axes are on a log scale. (b) The mean CPU time for completing the posterior probability calculation (Step 2) for a single set of demographic parameter values, given sampled coalescent trees from multiple loci. Both axes are on a log scale.

### Demographic model inference with and without ghost population

Once a sample of coalescent trees has been obtained, it can be used for analyses under multiple different demographic models, without having to resort to additional MCMC simulations. To demonstrate this we simulated 20 data sets, each with 50 loci and 4 gene copies from a single population which shares migrants with another unsampled population, a so-called “ghost” population. The sampled and unsampled populations occur in an IM model (Figure 1a) with parameters, *θ*_1_ = 1 (sampled population), *θ*_2_ = 5 (ghost), *θ*_*a*_ = 3, *m*_1_ = 2, *m*_2_ = 0.4 and *T*_*S*_ = 4. In step 1, 10,000 coalescent trees for each locus were sampled after 100,000 burn-in iterations and 100 thinning iterations from MCMC simulation. In step 2, the same set of coalescent trees was used repeatedly to infer two evolutionary scenarios. One is the data sampled from a single population that has not shared migrants with other populations. The other is an IM model in Figure 1a where the population of size *θ*_2_ was considered as “ghost”.

When a single population was assumed, its population size estimation was 4.15 which is much larger than the true size 1 (Figure 5). When an IM model was inferred, the estimations of all parameters were accurate (Figure 5). In particular, the estimated sampled population size was 1.338 close to the true value. Since we do not have a sample directly from the ghost population, the standard error for the ghost population size is large, but the confidence interval contains the true value. For model comparison, Akaike’s information criterion (AIC) was used. On 18 out of 20 replicates, IM model was selected rather than a single population model (Supplementary Table 3).

### False positives of likelihood ratio tests

The new method maximizes the joint posterior distribution, which is proportional to the joint likelihood when the prior distribution on demographic parameters is constant. Thus when working with uniform priors, and given a single sample of coalescent trees in step 1, the method can compare the maximum joint likelihoods, *L*_0_ and *L*_1_, under full and nested models, respectively (corresponding to alternative and null hypotheses, respectively) (see also Hey and Nielsen, 2007; Nielsen and Wakeley, 2001).

Recently a widely used method (implemented in IMa2) for LRTs for nested IM model comparisons (Hey and Nielsen, 2007) was shown to exhibit high false positive rates when actual divergence is low and the amount of data is not large (Cruickshank and Hahn, 2014). The cause of the high false positive rate was later shown to be largely due to the LRT being based on a marginal density that was not joint with the splitting time parameter and population sizes in the IM model. Hey *et al.* (2015) were able to generate a fully joint surface for a reduced model of 3 parameters, and showed that the observed distribution of the LRT test statistic followed the asymptotic distribution much more closely, and that the high false positive rate was much closer to target rate. Thus, because our new method estimates a joint posterior density in all demographic parameters, we were particularly interested in its LRT performance under the *small data, low divergence* context that exhibited high false positive rates for Cruickshank and Hahn (2014). We simulated 2, 10, 100, and 1,000 loci of 2 gene copies from each of two populations under recently diverged isolation models with *θ*_1_ = *θ*_2_ = *θ*_*a*_ = 5 and *T*_*S*_ = 0.5 or 10^*−*6^. We considered two low values for *T*_*S*_, including a value of effectively zero,*T*_*S*_ = 10^*−*6^, and a value of *T*_*S*_ = 0.5 which was used by Cruickshank and Hahn (2014) and Hey *et al.* (2015). For each case we simulated 100 replicates. In step 1, 1,000 coalescent trees for each locus were sampled after 100,000 burn-in and 100 thinning iterations. In step 2, the joint likelihoods are maximized under isolation model with same population sizes (null model) and IM model with same population sizes and same migration rates (alternative model) using the same set of trees. We computed LRT statistic *−*2(log*L*_0_ *−*log*L*_1_) for each case. The difference in the number of parameters between two models is 1.

**FIG. 5.**
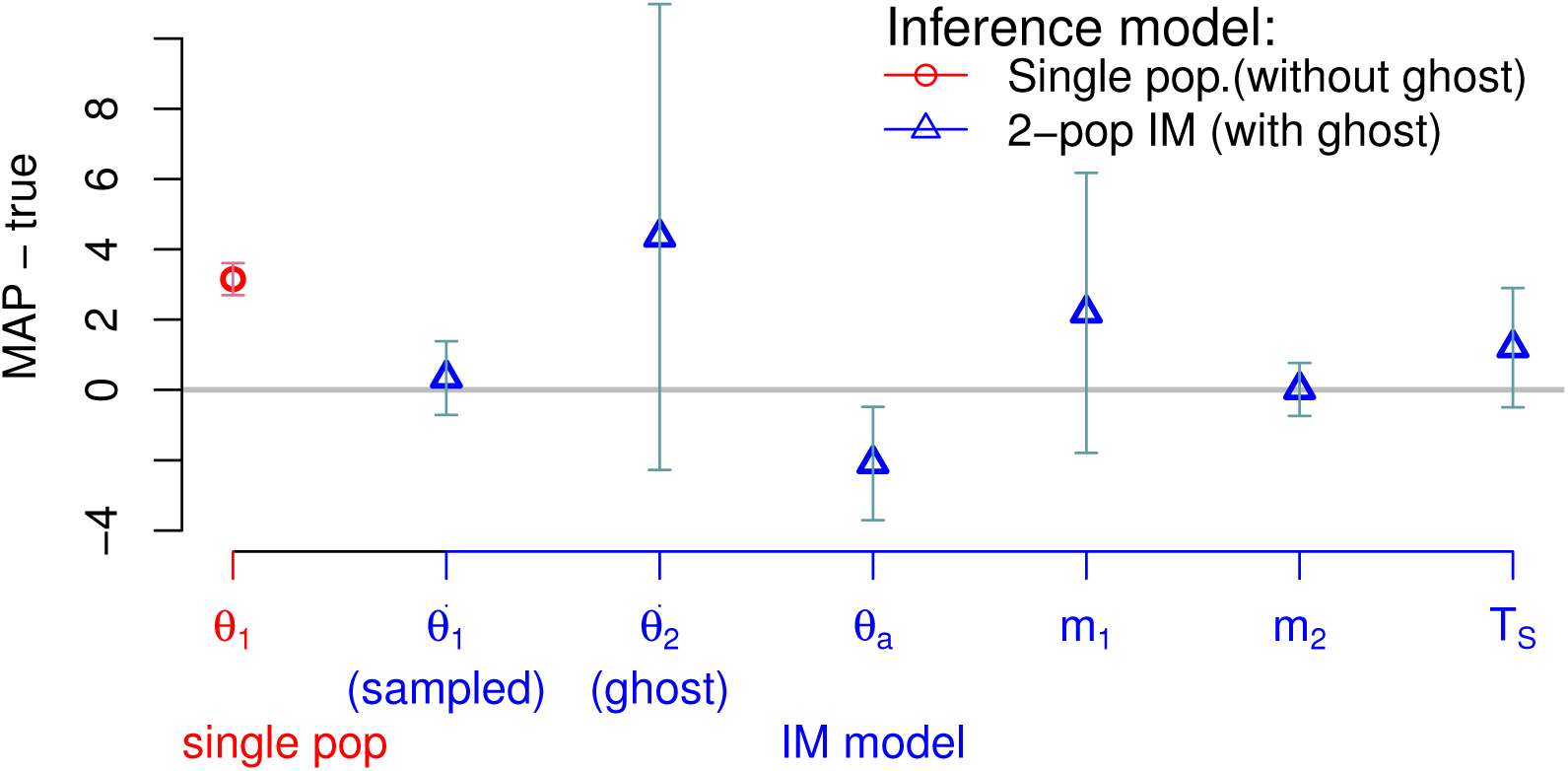
Estimation of demographic model with and without ghost population. The true simulation model is a 2-population IM model (*θ*_1_ = 1, *θ*_2_ = 5, *θ*_*a*_ = 3, *m*_1_ = 2, *m*_2_ = 0.4, *T*_*S*_ = 4) and we simulated DNA sequences from the population of size *θ*1. That is, the other population of size *θ*_2_ is a “ghost” population. Two demographic models were estimated: a single population model (∘, red) and IM model with ghost population (Δ, blue). Symbols represent the average difference over 20 replicates between MAPs and the true value. Bars represent the standard errors of parameter estimations.

**Table 2.** False positive rates of LRTs for migration rate are computed when the mixture distribution or an original 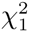 are considered as a null distribution. The proportions of zero values of LRTs are computed as well. The true simulation model is the 2-population isolation models with *θ*_1_ = *θ*_2_ = *θ*_*a*_ = 5 and *T*_*S*_ = 10*−*6, respectively. The number of loci varies from 2 to 1,000, and two gene copies are simulated from each population.

| No. loci | 2 | 10 | 100 | 1,000 |
| --- | --- | --- | --- | --- |
| False positive rate (mixture) | 0.01 | 0.05 | 0.11 | 0.02 |
| False positive rate ($\chi_1^2$) | 0.01 | 0.02 | 0.05 | 0.01 |
| Proportion of zero LRTs | 0.25 | 0.12 | 0.34 | 0.52 |

**Table 3.** False positive rates of LRTs for migration rate are computed when the mixture distribution or an original 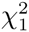 are considered as a null distribution. The proportions of zero values of LRTs are computed as well. The true simulation model is the 2-population isolation model with *θ*_1_ = *θ*_2_ = *θ*_*a*_ = 5 and *T*_*S*_ = 0.5. The number of loci varies from 2 to 1,000, and two gene copies are simulated from each population.

| No. loci | 2 | 10 | 100 | 1,000 |
| --- | --- | --- | --- | --- |
| False positive rate (mixture) | 0.03 | 0.14 | 0.03 | 0.13 |
| False positive rate ($\chi_1^2$) | 0.01 | 0.06 | 0.03 | 0.10 |
| Proportion of zero LRTs | 0.34 | 0.29 | 0.33 | 0.28 |

Typically when comparing two models that differ by one parameter the appropriate asymptotic distribution of LRT statistic is the *χ*^2^-distribution with 1 degree of freedom. However for the present case of the true parameter value equal to zero and on the boundary of the parameter space, the asymptotic distribution is a mixture distribution of zero with probability 0.5 and 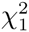 with probability 0.5 (Chernoff, 1954; Self and Liang, 1987). That is, we expect a half of LRTs to be zero when *m*_1_ =*m*_2_ =0. Therefore, we examined the proportion of zero LRTs and the false positive rates using two critical values, 2.705 and 3.841, from the mixture and original 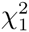 distributions with significance level 5%.

Table 2 shows the false positive rates and the proportion of zero values of LRTs. When the true splitting time is near zero, *T*_*S*_ = 10^*−*6^ the results show a false positive rate close to the expected rate. On 1,000-locus data sets, the LRT statistic seems to follow the mixture distribution (Supplementary Figure 6): the false positive rate is 2% and 52% of data sets have zero LRTs. In this case, the original test with 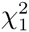 distribution shows conservative results. While the false positive rate on 100-locus case is elevated, those on smaller data sets are 5% or less. The proportions of zero LRTs on 100-locus or smaller data sets are 12%-34%, lower than the expected proportion of 50%. When the true model has a splitting time of *T*_*S*_ = 0.5, the MAPs of the parameters under isolation model and IM model, respectively, come closer to the true values with more loci (Supplementary Table 5). However, we observed some elevation of false positives (Table 3), though much smaller than when *T*_*S*_ is not in the joint distribution (Cruickshank and Hahn, 2014), with LRT values seeming to depart from the mixture distribution which is the limiting distribution of LRT when the number of loci goes to the infinity (Supplementary Figure 6). The false positive rates on 2-locus and 100-locus cases are smaller than 5% but larger than 5% on 10-locus and 1000- locus cases. The proportions of zero LRTs range from 28% to 34%. When the null hypothesis is rejected, migration rate and splitting time are always overestimated and the population size is underestimated (Supplementary Figure 5). This pattern indicates that the divergence of sequences under IM models with such large splitting time and migration rate is similar to that of the simulation model of zero migration rate and small splitting time.

### Evolutionary history of western and central common chimpanzees

We applied the new method to the demographic history of two common chimpanzee subspecies, *Pan troglodytes (P. t.) troglodytes* from Central Africa and *P. t. verus* from West Africa. These subspecies have been studied previously using IM models with small numbers of loci (Becquet and Przeworski, 2007; Hey, 2010a; Won and Hey, 2005). These and other studies (Caswell *et al.*, 2008; Wegmann and Excoffier, 2010) reported finding a signal of gene exchange between the subspecies with a divergence times of several hundred thousand years.

We aligned three sequences from each of two subspecies from the great ape genome project (Prado-Martinez *et al.*, 2013) by using the human genome reference (version 18). We partitioned the whole genome into non-overlapping segments of size 10,000 bps and selected 1,000 segments at random. In order to minimize a potential influence from recombination within a locus, each segment was separated into haplotype blocks using the four-gamete criterion (Hudson and Kaplan, 1985) and one block was selected at random from each segment. The average length of 1,000 loci was 4,206 base pairs. In step 1 of the analysis, an improper prior was assumed and 3,000 coalescent trees, scaled by per-site mutation rate, were sampled every 100 iterations after a burn-in of 100,000 iterations for each locus. Several MCMC diagnostics was carried out to ensure convergence (Supplementary Note and Supplementary Fig 3-4). In step 2, we estimated three population sizes of each of *P. t. troglodytes* and *P. t. verus* and their common ancestor, two migration rates and divergence time of them. The upper bounds for population sizes, divergence time and migration rates were 0.1, 0.01 and 1000, respectively. We used 48 CPUs for the step 1 analysis and it took around 3 hours. The step 2 analysis took around 17 hours on 196 CPUs.

Table 4 shows the parameter estimates obtained with our new method using 1,000 loci together with estimates from previous studies that used an IM model. These include Won and Hey (2005) and Hey (2010a) who used a 6-parameter IM model with 48 and 73 loci, respectively, and Becquet & Przeworski (2007) who analyzed 68 loci using a 5-parameter IM model with a single symmetric migration rate. All of these studies are broadly consistent with each other and suggest a model in which *P. t. troglodytes* is estimated to be about 4 times larger than that *θ*_2_ of *P.t. verus*, with gene flow occurring since their separation several hundred thousand years ago.

Our new estimate of the migration rate *m*_1_ from *P. t. verus* to *P. t. troglodytes* is larger than previously reported (2*N*_1_*m*_1_ = 0.878), but our estimate of *m*_2_ for the opposite direction is very close to zero (2*N*_2_*m*_2_ = 3.113*e*-13), which is consistent with Won and Hey (2005). In contrast Hey (2010a) reported a significant migration rate *m*_2_ under a 2-population IM model and Becquet & Przeworski (2007) reported a bidirectional rate of 2*N*_2_*m* = 0.1575 where *m* = *m*_1_ = *m*_2_. Overall the estimated model under the new method is consistent with results from previous studies. It is useful to note that all of these previous studies generated estimates from the marginal posterior distributions, whereas our analysis using MIST provides an estimate based on the full joint posterior density.

## Discussion

MCMC-based Isolation with Migration analyses have come to play a critical role in the analysis of population structure and of recent speciation events (Gronau *et al.*, 2011; Hey and Pinho, 2012; Payseur and Rieseberg, 2016; Pinho and Hey, 2010; Schraiber and Akey, 2015). The innovations presented here will enable the inclusion of larger portions of the genome, and provide a path for studying a wider range of demographic and phylogenetic models.

A major roadblock for existing MCMC based approaches that allow for extensive gene flow and population splitting is the non-independence of loci in demographic models with multiple time epochs. It is the updating of these epochs (e.g. splitting time *T*_*s*_) that must be done jointly for all loci, and that causes low acceptance rates in the Markov chain simulation when there are large numbers of loci (Wang and Hey, 2010). By using importance sampling of coalescent trees and removing the demographic model from the MCMC phase of the study, the MCMC update and sampling processes in the new method can treat loci independently.

Another major hurdle for MCMC-based methods that include migration over wide time periods are the complexities and time required to appropriately update genealogies that include migration paths. Our new approach includes an exact accounting of all possible migration path histories in the second phase of the analysis, allowing for the removal of migration paths from the MCMC phase and allowing for the importance sampling approach that does not rely upon a demographic model.

With a simpler Markov chain simulation that treats loci independently, we have no need for the use of metropolis-coupling (Geyer, 1991) or parallel tempering methods (Swendsen and Wang, 1986) that rely upon running multiple heated chains. In our experience a single Markov chain simulation for as many as 10,000 loci proceeds smoothly without mixing difficulty.

The emphasis on problems with large numbers of loci and small numbers of gene copies per locus is appropriate for many demographic problems, for which the optimal sampling effort favors more loci over more gene copies per locus (Cruickshank and Hahn, 2014; Felsenstein, 2006; Hey, 2010b; Hey *et al.*, 2015). The new method is designed to scale well with the number of loci, but limited to low numbers of gene copies per locus because the computing times and required memory size grows exponentially with the number of gene copies per locus (Supplementary Table 4). In the calculation of the posterior probability in the second step of the analyses, the new method employs transition rate matrices that are constructed for the unique ranked tree topologies with *population* labels among the sampled coalescent trees. The number of ranked tree topologies exponentially increases with the number of gene copies (Semple and Steel, 2003) and the sizes of transition rate matrices for each ranked tree topology are exponentially growing with the number of gene copies as well (Andersen *et al.*, 2014). For example, there are 7,248 unique ranked tree topologies on 8 gene copies, while 7 unique ranked tree topologies on 4 gene copies. The CPU times and physical memory usages in step 2 rapidly increased from 4 to 8 gene copies (Supplementary Table 4).

**Table 4.** Estimation of demography for two chimpanzee subspecies. Maximum a *posteriori* estimates (in parenthesis) of demography for *P. t. troglodytes* and *P. t. verus* from 3,000 loci of 6 sequences. Model parameter estimates are shown on a demographic scale, using a per-site mutation rate per generation of 2 × 10^*−*8^ and assuming 20 years per generation, and scaled by mutation rates in parentheses. Demographic units are individuals (*θ*), migration rate per gene copy per generation (*m*), and years *T*_*S*_.

| Method | No. loci | $\theta_1$ | $\theta_2$ | $\theta_a$ | $m_1$ | $m_2$ | $T_S$ |
| --- | --- | --- | --- | --- | --- | --- | --- |
| New method | 1,000 | 27,081.38<br>(0.00217) | 6,342.3<br>(0.00051) | 10,808.71<br>(0.00086) | 1.621e-05<br>(810.434) | 5.147e-18<br>(2.573e-10) | 347,732<br>(0.00035) |
| Won & Hey (2005)<sup>a</sup> | 50 | 27,900 | 7,600 | 5,300 | 9.2115e-6 | 1.1842e-7 | 422,000 |
| Hey (2010)<sup>b</sup> | 73 | 27,832.67 | 7191.48 | 8,399.21 | 9.108e-6 | 7.0496e-6 | 410,000 |
| Becquet *et al.* (2007)<sup>c</sup> | 68 | 33,000 | 9,750 | 15,000 | - | - | 439,000 |

<sup>a-b</sup> IMa2(Hey and Nielsen, 2007) was applied; <sup>a</sup> Geometric mean of mutation rate per locus per generation of 7.808e-6 and 15 years per generation were assumed; <sup>b</sup> Geometric mean of mutation rate per locus of 9.108e-6 and 20 years per generation were used; <sup>c</sup> MIMAR (Becquet and Przeworski, 2007) was applied. The per-site mutation rate per generation of $2 \times 10^{-8}$ and 20 years per generation assumed.

The new method is also unique in efficiently providing the joint MAP estimation of any given demographic model and for allowing model comparisons based on joint MAP estimates obtained under different models. Since typical Bayesian methods simulate demographic parameters and genealogies from an MCMC simulation, a very large number of samples is needed to find joint estimation requires. IMa2 cannot provide the joint estimation, since it simulates splitting times from an MCMC simulation and estimates the marginal or partial joint distribution for migration rates. The LRTs comparing marginal distributions gave rise to high false positive rates. The new method separate the incorporating processes of migration paths and coalescent trees and it enables to estimate the joint distribution of all parameters from a set of coalescent trees. LRTs using the new method compare joint distributions of all parameter estimations rather than marginal or partial joint distributions, and it solved the high false positive problem of LRTs for migration rates. On the small data sets, however, the distribution of the LRT statistic seemed to have a heavier tail than the null distribution which is the limiting distribution as the number of loci goes to the infinity. This departure from the limiting distribution led to more false positives than expected and the reliability of LRT seems to be affected by a small sample size. More studies are needed to discover how large sample is required for LRTs and to ensure possible other violation of the assumptions underlying LRTs.

## Acknowledgments

We thank Vitor Sousa for helpful discussion on an early version of the new method and Arun Sethuraman for writing a part of codes of MIST. This research was support under NIH grant R01GM078204 to J. Hey.

## References

Andersen, L. N., Mailund, T., and Hobolth, A. 2014. Efficient computation in the im model. Journal of Mathematical Biology, 68(6): 1423–1451.

Asmussen, S. 2003. Applied Probability and Queues. Springer-Verlag, New York, NY.

Bahlo, M. and Griffiths, R. C. 2000. Inference from gene trees in a subdivided population. Theoretical Population Biology, 57(2): 79–95. Article 306EQ THEOR POP BIOL.

Becquet, C. and Przeworski, M. 2007. A new approach to estimate parameters of speciation models with application to apes. Genome Research, 17(10): 1505–1519.

Beerli, P. and Felsenstein, J. 1999. Maximum-likelihood estimation of migration rates and effective population numbers in two populations using a coalescent approach. Genetics, 152(2): 763–73.

Berner, D., Grandchamp, A.-C., and Hendry, A. P. 2009. Variable progress toward ecological speciation in parapatry: Stickleback across eight lake-stream transitions. Evolution, 63(7): 1740–1753.

Caswell, J. L., Mallick, S., Richter, D. J., Neubauer, J., Schirmer, C., Gnerre, S., and Reich, D. 2008. Analysis of chimpanzee history based on genome sequence alignments. PLoS Genetics, 4(4).

Chernoff, H. 1954. On the distribution of the likelihood ratio. Ann. Math. Statist., 25(3): 573–578.

Cong, Q., Borek, D., Otwinowski, Z., and Grishin, N. V. 2015. Tiger swallowtail genome reveals mechanisms for speciation and caterpillar chemical defense. Cell Reports, 10(6): 910 – 919.

Cruickshank, T. E. and Hahn, M. W. 2014. Reanalysis suggests that genomic islands of speciation are due to reduced diversity, not reduced gene flow. Molecular Ecology, 23(13): 3133–3157.

Felsenstein, J. 1988. Phylogenies from molecular sequences: Inference and reliability. Annual Review of Genetics, 22(1): 521–565.

Felsenstein, J. 2006. Accuracy of coalescent likelihood estimates: Do we need more sites, more sequences, or more loci? Molecular Biology and Evolution, 23(3): 691–700.

Geraldes, A., Basset, P., Gibson, B., Smith, K. L., Harr, B., Yu, H.-T., Bulatova, N., Ziv, Y., and Nachman, M. W. 2008. Inferring the history of speciation in house mice from autosomal, x-linked, y-linked and mitochondrial genes. Molecular Ecology, 17(24): 5349–5363.

Geyer, C. J. 1991. Markov chain monte carlo maximum likelihood. Computing Science and Statistics, Proceedings of the 23rd Symposium on the Interface, pages 156–163.

Griffiths, R. C. 1989. Genealogical-tree probabilities in the infinitely-many-site model. Journal of Mathematical Biology, 27(6): 667–680. have pdf Perhaps the first paper to explicitly integrate over genealogies, though Felsenstein pointed out the general approach (albeit not via recursion) in his 1988 review.

Griffiths, R. C. and Tavaré, S. 1994. Simulating probability distributions in the coalescent. Theoretical Population Biology, 46: 131–159. finite site algorithms; xrx,.

Gronau, I., Hubisz, M., Gulko, B., Danko, C., and Siepel, A. 2011. Bayesian inference of ancient human demography from individual genome sequences. Nature Genetics, 43(10): 1031–1034.

Hey, J. 2010a. The divergence of chimpanzee species and subspecies as revealed in multipopulation isolation-with-migration analyses. Molecular Biology and Evolution, 27(4): 921–933.

Hey, J. 2010b. Isolation with migration models for more than two populations. Molecular Biology and Evolution, 27(4): 905–920.

Hey, J. and Nielsen, R. 2004. Multilocus methods for estimating population sizes, migration rates and divergence time, with applications to the divergence of drosophila pseudoobscura and d. persimilis. Genetics, 167(2): 747–760.

Hey, J. and Nielsen, R. 2007. Integration within the felsenstein equation for improved markov chain monte carlo methods in population genetics. PNAS, 104(8): 2785–2790.

Hey, J. and Pinho, C. 2012. Population genetics and objectivity in species diagnosis. Evolution, 66(5): 1413–1429.

Hey, J., Chung, Y., and Sethuraman, A. 2015. On the occurrence of false positives in tests of migration under an isolation-with-migration model. Molecular Ecology, 24(20): 5078–5083.

Hobolth, A., Andersen, L. N., and Mailund, T. 2011. On computing the coalescence time density in an isolation-with-migration model with few samples. Genetics, 187(4): 1241–1243.

Hudson, R. R. 2002. Generating samples under a wrightfisher neutral model of genetic variation. Bioinformatics, 18(2): 337–338.

Hudson, R. R. and Kaplan, N. L. 1985. Statistical properties of the number of recombination events in the history of a sample of dna sequences. Genetics, 111(1): 147–164.

Kimura, M. 1969. The number of heterozygous nucleotide sites maintained in a finite population due to steady flux of mutations. Genetics, 61: 893–903.

Kuhner, M. K. 2008. coalescent genealogy samplers: windows into population history. Trends in Ecology & Evolution, 24(2): 86–93.

Kuhner, M. K., Yamato, J., and Felsenstein, J. 1995. Estimating effective population size and mutation rate from sequence data using metropolis-hastings sampling. Genetics, 140(4): 1421–30.

Lopes, J. S., Balding, D., and Beaumont, M. A. 2009. Popabc: a program to infer historical demographic parameters. Bioinformatics, 25(20): 2747–2749.

Moodley, Y., Linz, B., Yamaoka, Y., Windsor, H. M., Breurec, S., Wu, J.-Y., Maady, A., Bernh¨oft, S., Thiberge, J.-M., Phuanukoonnon, S., Jobb, G., Siba, P., Graham, D. Y., Marshall, B. J., and Achtman, M. 2009. The peopling of the pacific from a bacterial perspective. Science, 323(5913): 527–530.

Nielsen, R. 2000. Estimation of population parameters and recombination rates from single nucleotide polymorphisms. Genetics, 154(2): 931–942.

Nielsen, R. and Wakeley, J. 2001. Distinguishing migration from isolation: A markov chain monte carlo approach. Genetics, 158(2): 885–896.

Payseur, B. A. and Rieseberg, L. H. 2016. A genomic perspective on hybridization and speciation. Molecular Ecology.

Pinho, C. and Hey, J. 2010. Divergence with gene flow: Models and data. Annual Review of Ecology, Evolution, and Systematics, 41(1): 215–230.

Prado-Martinez, J., Sudmant, P. H., Kidd, J. M., Li, H., Kelley, J. L., Lorente-Galdos, B., Veeramah, K. R., Woerner, A. E., O/’Connor, T. D., Santpere, G., Cagan, A., Theunert, C., Casals, F., Laayouni, H., Munch, K., Hobolth, A., Halager, A. E., Malig, M., Hernandez-Rodriguez, J., Hernando-Herraez, I., Prufer, K., Pybus, M., Johnstone, L., Lachmann, M., Alkan, C., Twigg, D., Petit, N., Baker, C., Hormozdiari, F., Fernandez-Callejo, M., Dabad, M., Wilson, M. L., Stevison, L., Camprubi, C., Carvalho, T., Ruiz-Herrera, A., Vives, L., Mele, M., Abello, T., Kondova, I., Bontrop, R. E., Pusey, A., Lankester, F., Kiyang, J. A., Bergl, R. A., Lonsdorf, E., Myers, S., Ventura, M., Gagneux, P., Comas, D., Siegismund, H., Blanc, J., Agueda-Calpena, L., Gut, M., Fulton, L., Tishkoff, S. A., Mullikin, J. C., Wilson, R. K., Gut, I. G., Gonder, M. K., Ryder, O. A., Hahn, B. H., Navarro, A., Akey, J. M., Bertranpetit, J., Reich, D., Mailund, T., Schierup, M. H., Hvilsom, C., Andres, A. M., Wall, J. D., Bustamante, C. D., Hammer, M. F., Eichler, E. E., and Marques-Bonet, T. 2013. Great ape genetic diversity and population history. Nature, 499: 471–475.

Price, K., Storn, R., and Lampinen, J. 2005. Differential Evolution: A Practical Approach to Global Optimization. Springer, New York.

Robert, C. P. and Casella, G. 2013. Monte Carlo Statistical Methods. Springer Science & Business Media.

Schraiber, J. G. and Akey, J. M. 2015. Methods and models for unravelling human evolutionary history. Nat Rev Genet, 16(12): 727–740.

Self, S. G. and Liang, K.-Y. 1987. Asymptotic properties of maximum likelihood estimators and likelihood ratio tests under nonstandard conditions. Journal of the American Statistical Association, 82(398): 605–610.

Semple, C. and Steel, M. 2003. Phylogenetics. Oxford University Press, New York, NY.

Swendsen, R. H. and Wang, J.-S. 1986. Replica monte carlo simulation of spin-glasses. Physical Review Letters, 57(21): 2607.

Wang, Y. and Hey, J. 2010. Estimating divergence parameters with small samples from a large number of loci. Genetics, 184(2): 363–379.

Wegmann, D. and Excoffier, L. 2010. Bayesian inference of the demographic history of chimpanzees. Molecular Biology and Evolution, 27(6): 1425–1435.

Wilson, I. J. and Balding, D. J. 1998. Genealogical inference from microsatellite data. Genetics, 150(1): 499–510.

Won, Y.-J. and Hey, J. 2005. Divergence population genetics of chimpanzees. Molecular Biology and Evolution, 22(2): 297–307.

Zhu, T. and Yang, Z. 2012. Maximum likelihood implementation of an isolation-with-migration model with three species for testing speciation with gene flow. Molecular Biology and Evolution, 29(10): 3131–3142.

